# Bioplastic Production from simulated 100% *in situ* Mars resources

**DOI:** 10.64898/2026.09.14.748363

**Authors:** Fatima R. Martin, Max G. Schubert, Harley Greene, Tom Pedersen, Nathan D. Hicks, Jordan E. Mancuso, Keren Isaev, Amy Spens, Jonathan Liu, Edward Sukarto, Una Nattermann, Devon A. Stork, Erika A. DeBenedictis

## Abstract

A sustained human presence on Mars requires local production of bulk materials, particularly polymers. Prior approaches to space biomanufacturing rely on human waste streams, Earth-sourced consumables, or complex chemical infrastructure, limiting their ability to scale. Here we demonstrate production of polyhydroxyalkanoate (PHA) bioplastic from simulated 100% martian resources: regolith-derived soluble nutrients, acetate electrochemically fixed from martian atmosphere. and water. We screened 16 candidate organisms for growth in a chemically defined Mars medium and identified *Cupriavidus necator* H16 and *Pseudomonas putida* KT2440 as promising chassis organisms. Adaptive laboratory evolution totaling more than 10^12^ cumulative cell divisions improved both species’ tolerance to high concentrations of acetate and leached regolith. The top C. *necator* evolved isolate, referred to as sPL.001, produced more than three-fold higher PHA titer under simulated Mars conditions compared to its parent strain. These results establish a path to polymer production on Mars where consumable mass is derived entirely from local resources. decoupling bulk material production from Earth supply chains.

## BIOMANUFACTURING IS AN EFFECTIVE WAY TO MANUFACTURE POLYMERS WITH IN SITU MARTIAN RESOURCES

Today, human space exploration is fundamentally limited by the feasibility of transporting the substantial quantity and diversity of materials required to sustain human life to other planets. A sustained, robust human presence on Mars will require manufacturing from local materials. a concept in space science known as *in situ* resource utilization (ISRU). Among material classes, polymers such as plastics, fibers, and films rank among the top sources of mass consumption projected for crewed space missions^1^. At sufficient scale. such polymers could serve as structural materials for greenhouses that transmit visible light. block UV, and maintain pressure differentials under Mars-relevant conditions^2^, enabling a virtuous cycle of expanding habitation and production capacity.

Researchers have proposed various approaches to polymer production from partially local resources, often framed as circular economy strategies. These include growing organisms in media supplemented with human waste^3–6^, and designing closed-loop biomanufacturing systems premised on recycling waste from Earth supply chains^7^. While these approaches represent important early steps, they share a fundamental limitation: their productivity is ultimately constrained by the flux of material arriving from Earth. They cannot scale indefinitely on Mars.

An alternative approach is to fix carbon and produce polymers through purely chemical or heavily supported biological routes. Thermochemical routes from CO_2_ to polymers (e.g., via syngas) require multi-stage, high-temperature infrastructure resembling a chemical refinery^8^. Gas fermentation routes, in which H_2_ and CO_2_ are fed directly to acetogens or other gas-fermenting organisms, reduce chemical complexity but still require specialized media formulation, water electrolysis, and gas solubilization^9,10^, with additional infrastructure if nitrogen must also be fixed industrially^11^. Photosynthesis avoids the need for electrochemical carbon fixation^3^, but photosynthetic conversion of sunlight to biomass is far less photon-efficient than photovoltaic-driven processes^12,13^, thus requiring vast insulated, transparent growth enclosures to capture sufficient light and maintain biotic processes. Each of these routes can, in principle, operate from local resources, but all impose infrastructure costs that scale with the number and difficulty of processing steps between raw martian inputs and the final product.

Here, we demonstrate a hybrid approach of electrochemical CO2 fixation to acetate using a compact electrolyzer^12,14^. followed by heterotrophic biological conversion of acetate into a polymer, supported by nutrients in martian regolith and the presence of oxygen. Defined Mars Media (DMM) simulates the result of solubilizing regolith in water, and supplies nitrogen, phosphorus, sulfur, and other inorganic nutrients, along with the stressors of elevated salinity, perchlorate, and heavy metals^15^. Molecular oxygen can be generated via solid oxide electrolysis as demonstrated *in* situ by the MOXIE instrument aboard Perseverance^16^. The complete abiotic infrastructure footprint thus reduces to four systems: water harvesting. regolith solubilization, CO,-to-acetate electrolysis, and 02 electrolysis **(Figure 1)**. This hybrid approach builds out the infrastructure to support biomanufacturing in a bioreactor brought to Mars. Which provides radiation protection and controls temperature and pressure. The use of heterotrophic organisms that require an organic carbon source to grow and cannot fix carbon from CO2 or sunlight alone limits their capacity to proliferate in the Martian environment if released, with benefits for planetary protection^42^. Given this process architecture, we need a heterotrophic organism that grows on acetate with regolith-leached nutrients and produces a useful biopolymer.

**Figure 1.**
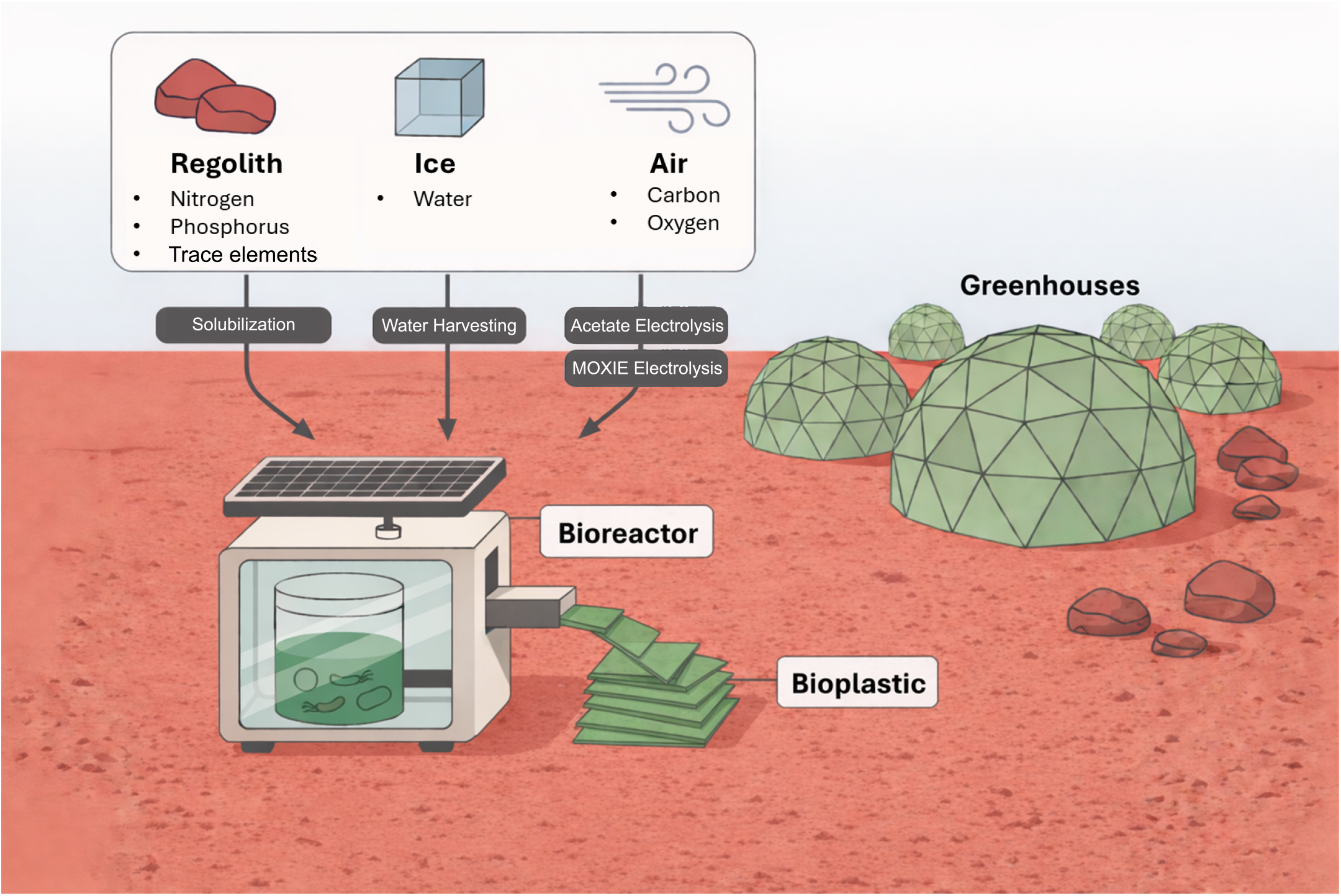
Minimal infrastructure for bioplastic production from Martian resources. A bioreactor deployed to Mars could support biomanufacturing microorganisms entirely on Mars-derived resources: inorganic nutrients from the regolith, water from ice. and oxygen and a carbon source derived from the atmosphere. Microbially produced bioplastic from these *in situ* resources could be used to construct greenhouses or for other building material needs.

We chose polyhydroxyalkanoates (PHAs) as our target polymer because many acetate-utilizing and oxygen-respiring organisms produce them natively, and they can be processed into 3D-printable filaments, flexible films, and structural fibers^17^. PHA copolymers have tunable mechanical properties ranging from rigid to elastomeric^18^, making them suitable for a wide range of applications, from greenhouse materials^19^ to airtight pressure vessels^20^, and they accumulate intracellularly as carbon storage granules, which can be extracted after cell harvest^21,22^ with minimal consumable usage^23^ or extruded directly^24^.

We therefore sought heterotrophic organisms capable of growth under simulated Martian conditions in DMM^15^, which contains Mars-relevant concentrations of nitrate and phosphate as essential inorganic nutrients. as well as the stressors of salinity, perchlorate, and heavy metals. We demonstrate and improve the production of PHAs to show the potential for scalable, fully *in situ* bioplastic production from entirely Martian resources.

## SCREENING CHASSIS ORGANISMS FOR GROWTH IN MARS CONDITIONS

Our proposed Mars-derived feedstock is a complete microbial growth environment, with no processing steps to remove toxins or enrich nutrients. While DMM contains fixed nitrogen as nitrate (N0_3_^-^), phosphorus as phosphate (Poi^-^). and other essential minerals, it simultaneously exposes organisms to elevated salinity, perchlorate (CI0_4_^-^), and metals. Acetate itself becomes inhibitory at high concentrations, as the protonated acid diffuses across membranes and disrupts cytoplasmic pH. A chassis organism for this biomanufacturing task must not merely survive this polyextreme environment but grow in high concentrations of it and have spare metabolic capacity to produce a desirable product.

We defined four physiological requirements for candidate organisms. First, it must use acetate as its sole carbon and energy source. Second. it must assimilate nitrate from regolith as its sole nitrogen source, since nitrogen in the Martian atmosphere is too dilute to support nitrogen fixation without enrichment^25^. Third, the organism must be prototrophic, or not require any nutrients other than those in DMM. as any complex supplements (amino acids, vitamins, etc.) cannot be sourced from raw Martian materials. Fourth, it must tolerate the combined stressor profile of leached regolith, at as high a concentration as possible. Additionally, we considered genetic tractability, or the availability of established transformation methods and characterized genetic parts, as a strong practical advantage for downstream strain improvement or adaptation to other products, though not as an absolute requirement.

We screened 16 organisms drawn from the astrobiology and industrial microbiology literature against the four physiological criteria described above **(Methods)**. Eight were excluded after failing one or more criteria in preliminary experiments. Oeinococcus *radiodurans*. despite its exceptional radiation tolerance. requires amino acid and vitamin supplementation and cannot grow prototrophically^26,27^. *Colwellia psychrerythraea*, a psychrophile, similarly cannot grow on unsupplemented defined medium^28^. *Vibrio* natriegens and Ha/omonas campaniensis did not grow in DMM, likely because DMM does not contain sufficient sodium for these halophiles before reaching toxic levels of perchlorate. *Klebsiella variicola* was unable to use nitrate, while *Xanthobacter autotrophicus* and Oech/oromonas *agitata* did not grow in DMM, and P/anococcus *halocryophilus* did not show prototrophic growth in our laboratory. We previously probed the growth of the remaining eight microbes using a series of DMM concentrations, reflecting the soluble minerals expected at different regolith-to-water ratios^15^ **(Figure 2A)**.

**Figure 2.**
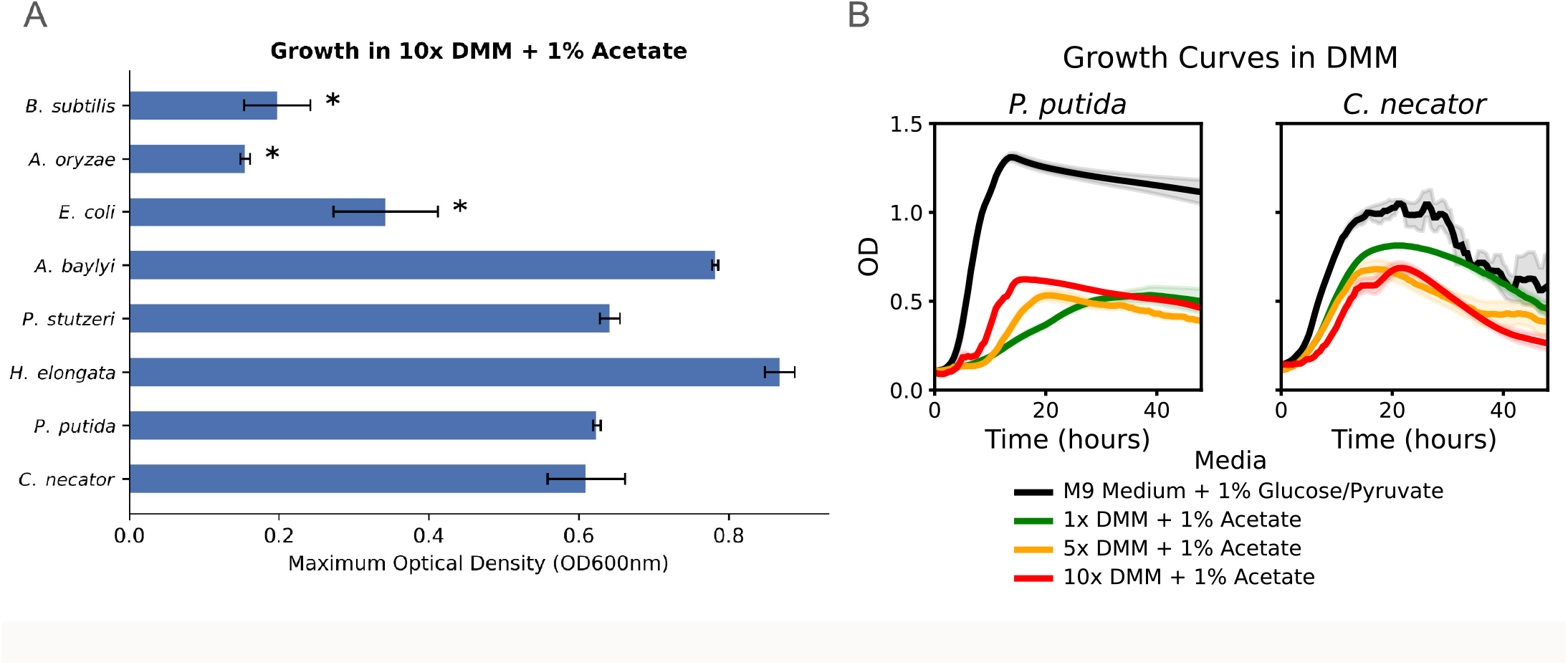
Organism screening for Mars bioprocess conditions. **A)** Maximum OD_600_ that bacterial cultures reached over 72 hour growth curves in a 10× formulation of DMM with 1% potassium acetate added (n=2). Asterisks reflect strains we identified in this low-volume assay that grew well in DMM initially but translated to no growth in DMM in larger volumes and over longer periods. See **Methods** for assay details. **B)** Growth curves (n=4) for *C. necator* and *P. putida*. Both strains grow best in permissive M9 medium (with 1% glucose for *P. putida* and potassium pyruvate for *C. necator* as a carbon source; black line), but are still able to grow robustly in DMM at 1× (green line), 5× (yellow line), and 10× (red line) concentrations with 1% potassium acetate. Shaded region indicates 95% confidence interval.

Two organisms emerged as the strongest candidates: Pseudomonas *putida* KT2440 and *Cupriavidus necator* H16. Both grew better than other screened organisms in the desired conditions, both are well-established biomanufacturing chassis with mature genetic toolkits. and both natively produce PHA. C. *necator* in particular is one of the best-characterized natural PHA producers^29^, and *P. putida* has a flexible metabolism that can produce medium-chain-length PHAs with distinct material properties^30^.

Both organisms grew substantially better in standard defined minimal media than in DMM. even at higher DMM concentrations that provide more total nutrients **(Figure 2B)**. This gap reflects a fundamental tension in our growth environment: higher leached regolith concentrations deliver more nutrients (nitrogen. phosphorus, trace elements) but also proportionally more stressors. At low DMM concentrations. strains grow reasonably well but are nutrient-limited; at higher concentrations, toxicity slows or halts growth. Improving tolerance to higher leached regolith concentrations would therefore simultaneously increase the nutrient supply available to the organism and improve bioproduction, a goal well-suited to adaptive evolution.

## ADAPTIVE LABORATORY MARS CONDITIONS EVOLUTION IN Mars conditions

In a previous pilot study. we used salt stress to compare adaptive laboratory evolution (ALE) against genome-wide saturating mutagenesis^31^ and functional genomics from extremophile organisms^32^. We concluded that ALE consistently captured the largest early fitness gains with the least upfront effort, making it the natural starting point for a complex, multi-stressor environment where the relevant genetic targets are not known in advance. We therefore performed ALE in both C. *necator* and *P. putida* in challenging concentrations of acetate and DMM. to improve tolerance to these stressors. thereby increasing the nutrient supply available for growth and bioproduction.

The ALE campaigns used serial passaging of triplicate 50 ml flask cultures over approximately six weeks. Weekly check-ins included 16S amplicon sequencing, to monitor for contamination. and inhibition assays. to track the populations’ increasing tolerance to the stressor. For C. *necator*, we began from an engineered parent strain derived from wild type (WT). containing a genomic integration which had modestly improved tolerance to DMM (unpublished data). and passaged cultures every 48 hours in 14x DMM with 1.25% potassium acetate for the first three weeks. then increased stressors to 20x DMM with 1.75% potassium acetate for the final three weeks. For *P. putida*, we began with the WT KT2440 isolate, initially passaging every 24 hours in 20x DMM with 1.5% potassium acetate, then shifting to 48-hour passages with 20x DMM in 2% potassium acetate to select for sustained growth over longer culture periods more representative of fed-batch bioproduction conditions. In both cases. we chose starting concentrations of stressors near the upper limit of what the parent strains could tolerate. then increased them as the populations adapted. Over the full course of the experiment we estimate more than 10^12^ cumulative cell divisions occurred per strain (2.5 x 10^12^ for *P. putida*. 10^12^ for C. necator). in line with the scale and duration of previous studies evolving strains to better tolerate stressors^33,34^.

Both species showed continual improvement in stress tolerance over the six-week ALE campaign **(Figure 3A**, additional details in **Methods)**. Populations progressively improved their ability to grow at higher DMM and acetate concentrations, indicating increasing fitness in the target environment. These improvements continued throughout the duration of the whole experiment, suggesting that additional improvements may be seen with further passaging.

**Figure 3.**
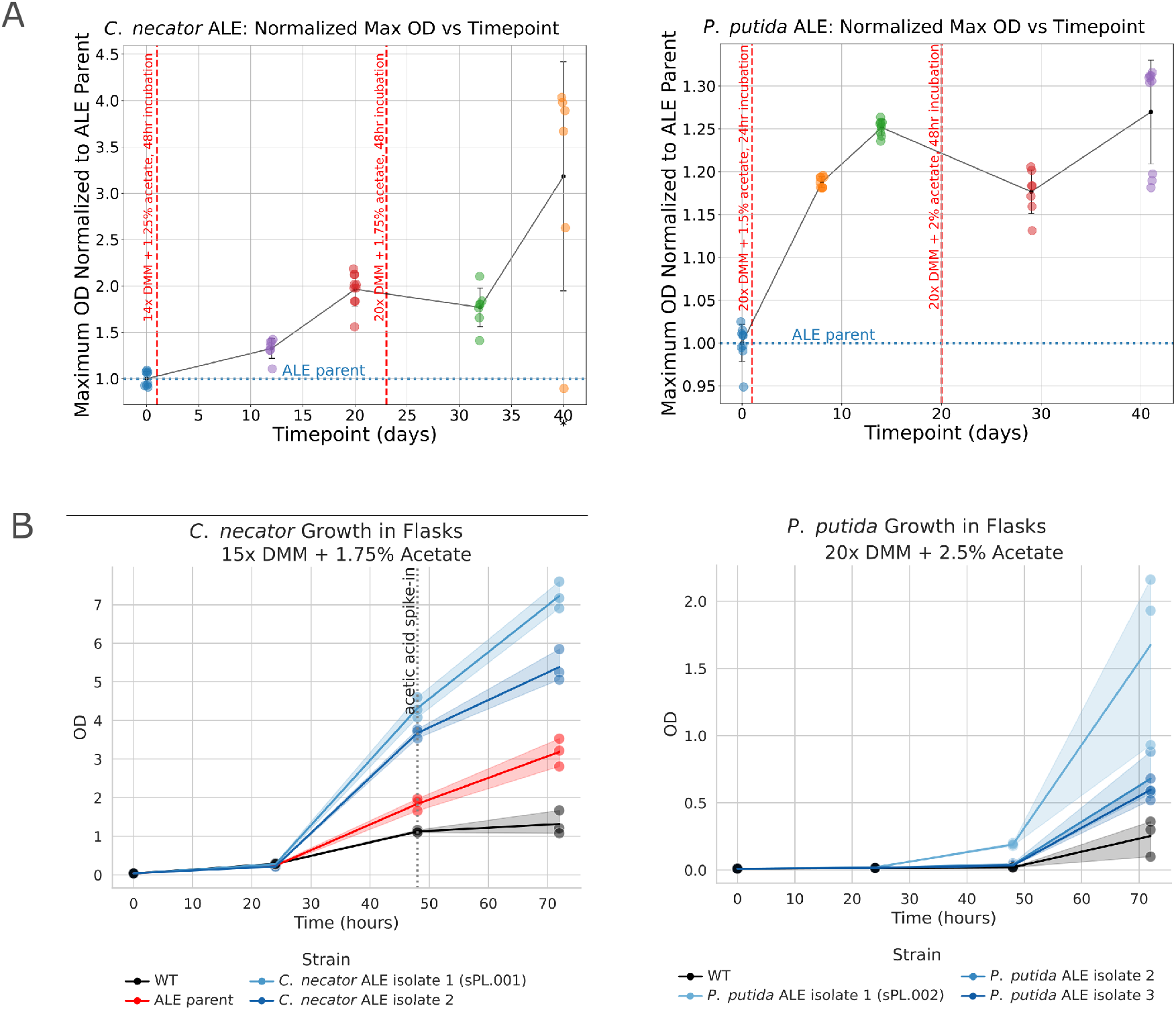
Adaptive evolution in Mars bioreactor-relevant conditions. A) Once weekly. we assessed the performance of our evolved populations using a growth curve assay in microtiter plates **(Methods).** Here those assessments are summarized by reporting the maximum growth curve OD_600_ achieved (n=3 for each flask). in the medium specified, normalized to ALE parent strain measurements for the species indicated. Error bars represent standard deviation between samples. B) Evolved C. *necator* and *P. putida* isolate performance in flask growth at 15x DMM + 1.75% acetate and 20x DMM + 2.5% acetate. respectively. across 72 hours (n=3). ALE isolates for C. *necator* and *P. putida* reached higher ODs compared to the wild type (WT) and ALE parent for *C. necator*. The top performing isolates (isolate 1 for both) were saved as sPL.OO1 and sPL.OO2. for *C. necator* and *P. putida* respectively. Shaded region indicates range.

From the final ALE populations, we isolated individual clones, performed whole-genome sequencing, and identified unique mutations relative to the parent strain. We tested tens of isolates per species by observing their growth curves across a range of DMM and acetate stress concentrations and selected strains that reached the highest biomass at the highest nutrient concentrations as top performers.

To assess whether improved low-volume growth in 96-well plates translated to a more realistic production context, we grew the three top isolates alongside their parent strains in 72-hour 1OO-ml shake flask cultures. For C. *necator*, we included periodic acetic acid spike-ins, to simulate a fed-batch process for pH management and carbon replenishment. *P. putida* showed higher tolerance to both DMM and acetate and was assessed in more challenging conditions, where acetic acid feeding was not necessary. Both protocols more closely resemble the operation of a bioreactor than low-volume batch growth in microtiter plates. The evolved isolates maintained their growth advantage over the parent strains in flasks **(Figure 3B)**, confirming that the ALE-derived tolerance gains are robust across culture conditions, culture density, and time scales relevant to bioproduction. The top performing isolates were glycerol stocked and hereafter referred to as sPL.O01 and sPL.OO2, for C. *necator* and *P. putida* respectively, for future work, such as forward planetary protection investigations^42^.

## EVOLVED MICROBES SHOW HIGHER BIOPRODUCTION IN MARS CONDITIONS

In biomanufacturing, improved growth does not always translate to improved product titer, as cell growth and product synthesis compete for shared precursors and energy, and adaptive mutations that improve fitness can redirect flux away from production pathways^35^. However, for intracellular storage products like PHAs the correlation is expected to be stronger because PHA accumulates as a carbon and energy reserve under normal growth conditions. We sought to confirm this directly by quantifying bioplastic production in our evolved isolates under Mars-relevant conditions. We observed evidence of PHA accumulation in both species, but focused further efforts on C. *necator* where the fraction of biomass accumulated as PHA is known to be highe.

We harvested the final C. *necator* biomass from the 72-hour 100 ml fed-batch flask culture experiments described previously. PHA was purified with a simple bleach lysis method^36^, and then washed, dried, and weighed. The best evolved isolate (sPL.001) produced over 1.5 g/L PHA, compared to under 0.5 g/L for the parent strain, a greater than three-fold improvement under identical simulated Mars conditions **(Figure 4A)**. This confirms that ALE-adapted strains are not merely surviving better in Mars-relevant conditions, but that they are far better suited for bioproduction on Mars.

**Figure 4.**
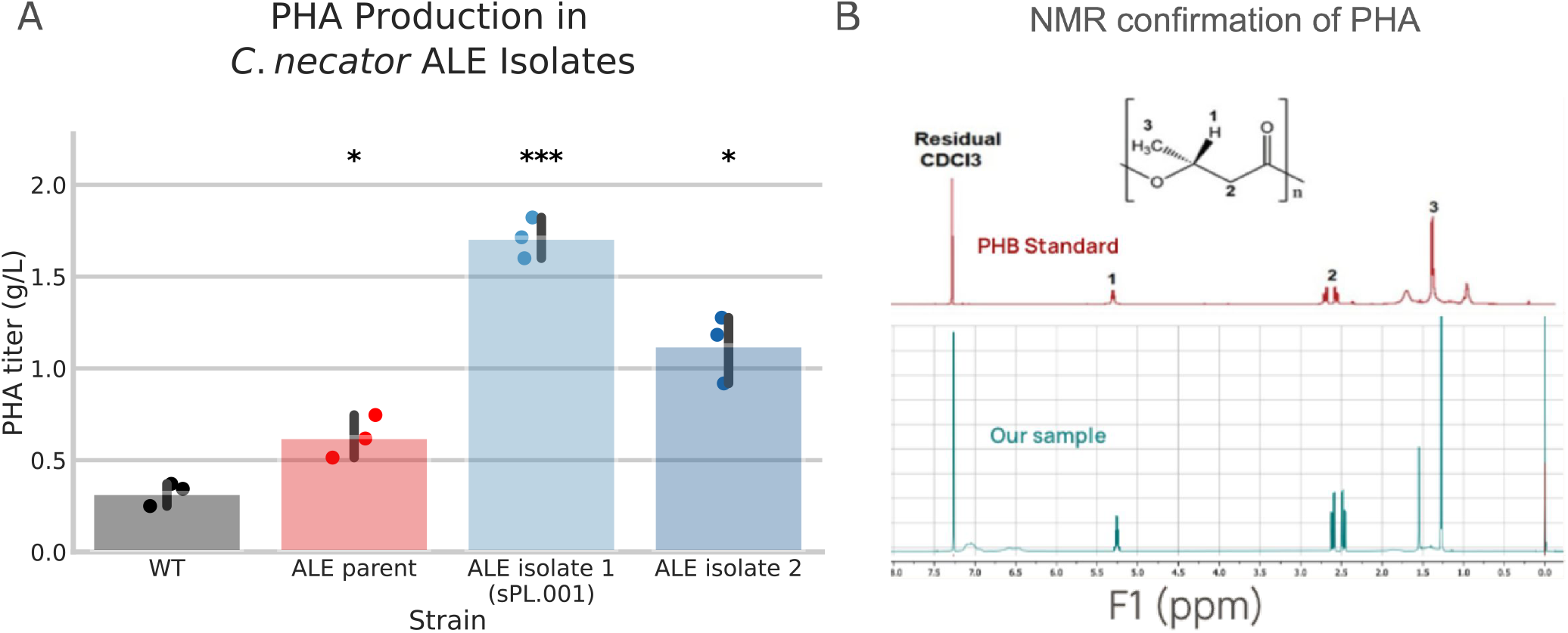
PHA bioproduction in Mars bioreactor conditions. A) PHA titer (g/L) for C. *necator* WT, ALE parent, and evolved isolates in 15x DMM + 1.75% potassium acetate (n=3). Two C. *necator* isolates make PHA at significantly higher titers than both WT and the ALE starting strain. This increase in production is due to evolved strains’ ability to grow to much higher ODs. Error bars depict the 95% confidence interval, and asterisks depict the results of a pairwise t-test against WT, with * depicting p < 0.05, and *** depicting p < 0.001. B) Proton nuclear magnetic resonance (^1^H-NMR) analysis of PHA, comparing a known PHB ^1^H-NMR spectrum^37^ (top, red) to PHB that we extracted from our evolved C. *necator* (blue, bottom). Key identifying peaks for PHB (labeled 1, 2, 3) are present in our sample. Extraneous peaks that appear in our sample at 0 ppm and 1.5 ppm are caused by TMS standard (internal control for NMR analysis) and water contamination, respectively.

To verify the identity of the purified material. we performed proton nuclear magnetic resonance CH-NMR) on the extracted polymer and compared the spectrum to a commercial standard of the PHA subtype polyhydroxybutyrate (PHB) produced by C. *necator*. Matching spectra confirmed that the material produced by our evolved C. *necator* in simulated Mars media and purified from cells in our simple protocol is the expected bioplastic polymer^37^ **(Figure 4B)**.

## CONCLUSIONS & FUTURE DIRECTIONS

These results demonstrate production of a useful polymer derived entirely from simulated Martian resources. No Earth-sourced consumables are required, meaning production capacity is unconstrained by upmass. It also does not rely on human waste for nutrients, making this process possible on Mars before human arrival, and can scale independent of human population size. It can be accomplished with a minimal infrastructure footprint: devices for regolith leaching: COrto-acetate electrolysis: a bioreactor: and access to power and 02, thus avoiding the footprint sprawl of more complex chemical refineries. Additionally, because the chassis organisms are heterotrophic and require an organic carbon source. they cannot proliferate in the Martian environment if released^42^. We show that ALE improved microbial tolerance to higher concentrations of leached regolith and supplemented acetate, and those tolerance gains translated directly to improved bioproduction. This is the first demonstration that adaptive evolution under simulated Martian conditions translates to improved biomanufacturing performance. Strains sPL.001 and sPL.002 represent our first efforts at designing microbes specifically to excel in Mars’ challenging chemical conditions.

sPL.001 produces over three-fold more PHB than its terrestrial parent strain in Mars-relevant conditions. A key open question is how close the yield of this process can approach optimized terrestrial PHB production. Our current measurements are from flask cultures and are not directly comparable to optimized fed-batch or continuous bioprocesses where productivity can reach 12 g/L/day^17.38^.

Much of this gap reflects inherently less optimal nutrients in DMM, including the fact of nitrate and acetate being more challenging sources of nitrogen and carbon, respectively. Some of this gap could be alleviated by further fermentation development, and some by further improving DMM tolerance. Tolerance had not plateaued by the end of our evolution campaigns, suggesting further adaptation would accrue more benefits. Equally important, the availability of evolved strains that grow robustly in Mars media now unblocks the development of Mars-specific bioreactor hardware: reactor geometry, feeding strategy, and process control can be optimized around an organism that actually performs in the target medium. How close to the terrestrial productivity maximum this combination of strain improvement and process engineering can achieve remains an open question.

Bioplastics are a diverse and tunable class of polymer whose material properties can be customized through biological engineering. PHA copolymers range from rigid to elastic^18^, and the two chassis microorganisms developed here (sPL.001 and sPL.002) produce complementary polymer types: C. necator accumulates short-chain-length PHAs, while *P. putida* produces medium-chain-length PHAs with distinct mechanical properties^30^ . These biomaterials can be used in diverse use cases as 3D-printable filaments for replacement parts. films, fibers, and potentially as structural materials for greenhouses^2,19^ used to construct enclosed agricultural or habitable regions on Mars from entirely Martian resources^39^. Further genetic engineering, downstream purification improvements^40^, biological containment studies, and ultimately flight testing could advance this process toward implementation.

## MATERIALS AND METHODS

### Media Formulation

Defined Mars Media (DMM) was formulated as previously reported^15^. Briefly, 1x DMM simulates the solution obtained from leaching 40 grams of martian regolith in 1 liter of water, drawing on data from the Phoenix Lander Wet Chemistry Laboratory, Curiosity Rover Sample Analysis at Mars mineralogical studies, and studies of synthetic regolith leachates. Similarly, 1x Mars Trace Micronutrients (MTM) represents the soluble trace metals fraction of this expected solution. Our inhibition curves in this study extend to very high concentrations of DMM (e.g. 30x DMM) in which we observed calcium sulfate (gypsum) precipitation, so we used a “low calcium” modification of DMM in which Ca^2^. is added at one tenth concentration for all experiments shown, allowing Ca^2^. to be soluble at the concentrations and conditions tested here.

### Screening Organisms For Growth In DMM + Acetate

Bacterial strains were obtained from the American Type Culture Collection (ATCC), Bacillus Genetic Stock Center (BGSC), Coli Genetic Stock Center (CGSC). and Leibniz Institute DSMZ-German Collection of Microorganisms and Cell Cultures (DSM) as indicated below. The full names of organisms included in Figure 2 are *Escherichia coli* MG1655 (CGSC 6300), Bacillus subti/is *PY79* (BGSC lA747). *Pseudomonas putida KT2440* (DSM 6125). *Pseudomonas stutzeri AW-1* (DSM 13592), *Halomonas* e/ongata *1H9* (ATCC 33173), *Cupriavidus necator H16* (DSM 428), *Azospira oryzae* PS (DSM 13638), and *Acinetobacter baylyi ADP1* (DSM 24193). Additional organisms screened include Deinococcus *radiodurans R1* (ATCC 13939), *Klebsiella variicola F2R9* (DSM 15968), *Vibrio natriegens PB111* (ATCC 14048), *Xanthobacter autotrophicus* 7C (DSM 432), *Colwellia psychrerythraea 34H* (ATCC BAA-681), *Dechloromonas* agitata *CKB* (DSM 13637), P/anococcus *halocryophilus Ori* (DSM 24743), and Ha/omonas campaniensis *5AG* (DSM 15293). Organisms were grown in their ATCC recommended medium and cryo-preserved in 25% glycerol at -80°C.

We prepared minimal media (M9) as described by Cold Spring Harbor Protocols^43^. Acetate was added from a stock solution of 20% w/v potassium acetate. The media used in Figure 2 are as follows: 1) M9 with 1% glucose and 1/100 ATCC Trace Mineral Supplement (ATCC-TMS), 2) 1x DMM with 1% potassium acetate, 50 mM N-ITris(hydroxymethyl)-methyl]-2-aminoethanesulfonic acid (TES, a common buffer), and 1x MTM, 3) 5x DMM with 1% potassium acetate, 50mM TES, and 5x MTM, 4) 10x DMM with 1% potassium acetate, 50 mM TES, and 10x MTM. For C. *necator*, in instances where 1% glucose was used for other strains, 1% potassium pyruvate was also added.

For benchmarking the strains in M9 and DMM media, we first started overnights of the strains in rich media: Luria Broth (LB) (most strains), Super Optimal Broth (SOB: Fisher Scientific 244310) (C. necator), LB (Fisher Scientific 244620) + 3% NaCl *(H*. e/ongata), or ALP medium41 minus lactate *(A*. oryzae). After 16 to 20 hours of growth at 30^°^C, the ODs of strains were measured. The benchmarking media was prepared as described above, and 100 µL was added to microtiter plates (Fisher Scientific 12-566-70). 5 µL of overnight culture normalized between 0.3 and 0.7 OD_600_ was inoculated into these media and growth curves were conducted for 72 hours at 30^°^C with high speed shaking and readings every 10 minutes, in either a BioTek Synergy HT or a BioTek Synergy Mx plate reader.

### ALE Passaging In Flasks

C. *necator* ALE cultures were grown in 250 ml baffled flasks with 50 ml of l4x DMM + 1.25% potassium acetate + 50 mM TES + 1/1000 ATCC-TMS for the first 3 weeks of ALE, then stressors were increased to 20x DMM + 1.75% potassium acetate + 50 mM TES + 1/1000 ATCC-TMS for the last 3 weeks of ALE. Kanamycin sulfate was also included at 25 µg/ ml in cultures to prevent contamination, as the C. *necator* ALE parent contained a genomically-integrated kanamycin resistance cassette. After cultures had incubated shaking at 30’C for 48 hours, 0D_600_ was measured, and 2 ml of saturated culture was passaged into fresh media.

Similarly, *P. putida* ALE cultures were grown in 250 ml baffled flasks with 50 ml of 20x DMM + 1.5% potassium acetate + 50 mM TES + 1/1000 ATCC-TMS with 24 hour incubation for the first 3 weeks of ALE. For each passage, 500 µL of saturated culture was inoculated into fresh media. Stressors were increased to 20x DMM + 2% potassium acetate + 50 mM TES + 1nooo ATCC-TMS with 48 hour incubation for the last 3 weeks of ALE.

For each week of passaging, we checked ALE populations to catch any contamination early and measure how much tolerance to stressors had improved. 16S PCR (forward primer AGAGTT TGATCCTGGCTCAG, reverse primer ACGGCTACCTTGTTACGACTT) of each culture was sequenced (Premium PCR, Plasmidsaurus), which allowed us to check for low levels of contamination in the population that Sanger sequencing would otherwise have missed. l ml from each culture was cryo-preserved at -80’C after adding glycerol to 25% volume. We also measured kinetic growth curves across a range of DMM and acetate concentrations for each population and the ALE parent to compare how growth phenotypes had changed between unevolved and evolved populations. Cumulative cell divisions are computed by assessing the number of cells at the end of each culture, subtracting the cells used to inoculate that culture, and summing across all cultures. The number of cells was estimated from OD_600_ by using the common conversion 8 x 10^8^ cells / ml x 0D_600_.

### Inhibition Curve Assay

To quantify relative fitness of strains in DMM and potassium acetate, we measured growth curves across a range of concentrations for DMM and acetate. Using the maximum OD_600_ that cultures reach over 72 hours, we can make an inhibition curve **(Figure 5)** that allows us to easily identify strains with the best tolerance for the stressor of interest. We typically assessed inhibition both across a range of DMM or acetate concentration while holding the other constant, rather than varying both types of stressors together. This allows us to conclude that evolved strains better tolerate both stressors independently.

**Figure 5.**
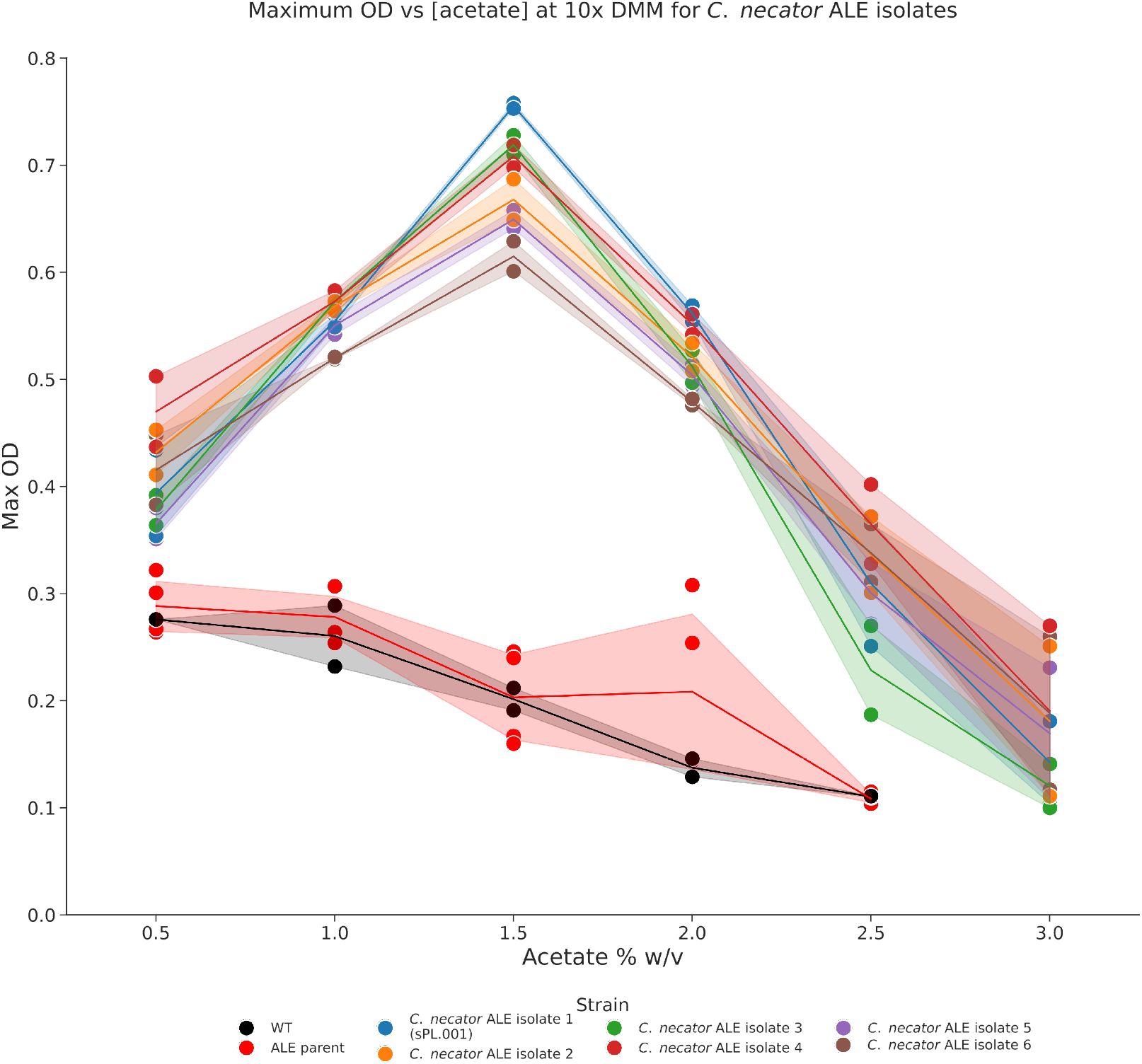
Representative acetate inhibition curve for ALE isolates. Evolved *C. necator* strains reach higher ODs at all concentrations of acetate and can tolerate higher concentrations of acetate (n=3). Notably, early in the inhibition curve, additional acetate results,n more growth by delivering more nutrients. but higher concentrations of acetate begin to inhibit growth. Shaded region indicates standard deviation.

For the inhibition curve assay. strains were revived from glycerol stocks and inoculated into SOB for overnight growth at 30^°^C. ODs of saturated cultures were measured in cuvettes and cultures were pelleted to remove the SOB supernatant to prevent any rich media carryover. Cultures were normalized in TES-buffered saline (10 mM TES + 1% w/v NaCl) to OD_600_ 0.7. To prepare the plate with a gradient of stressor concentrations. two media were made at the highest and lowest concentrations of the stressor of interest. while all other components and stressors were held constant. A Hamilton STARlet liquid handler was used to combine media at given volumes in a 96 deep-well plate to achieve intermediate concentrations. 100 µL of the resulting media was then inoculated with 5 µL of OD-normalized seed culture in each well. Growth curves were conducted for 72 hours at 30^°^C with high speed shaking and OD_600_ read every 10 minutes, in either a BioTek Synergy HT or a BioTek Synergy Mx plate reader **(Figure 6)**.

**Figure 6.**
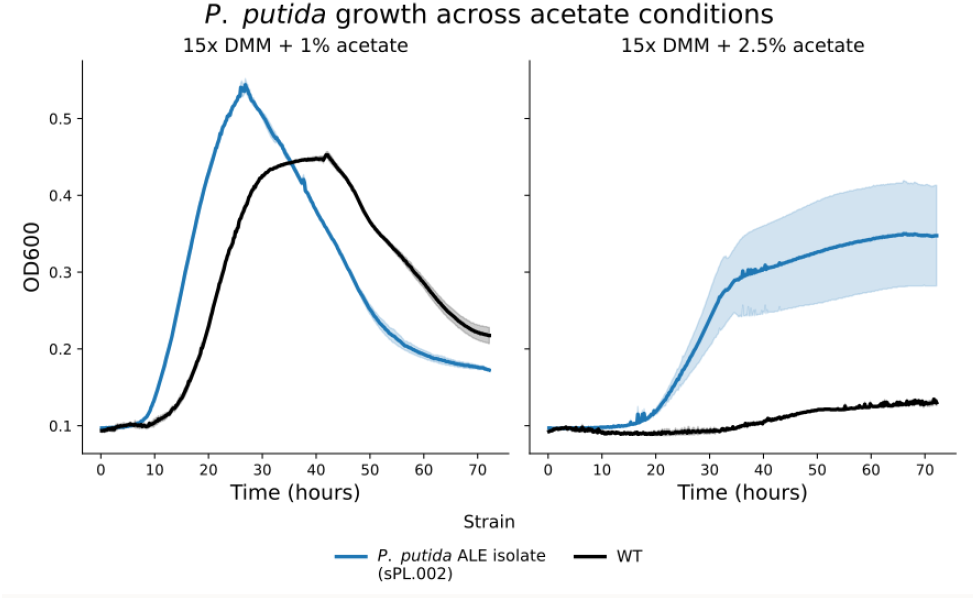
Growth across acetate conditions. *P. putida* growth curves at 15x DMM + 1% acetate and 15x DMM + 2.5% acetate (n=3). At 1% acetate, cultures experience a steep death phase due to low available acetate. At 2.5% acetate, OD is much more stable, due to excess acetate. Shaded region indicates 95% confidence interval.

### Obtaining, Sequencing, And Testing Isolates

To obtain isolates from ALE populations, glycerol stocks from final ALE timepoint populations were struck out on agar plates, either SOB solidified with 1.5% agar or 15x DMM + 1% potassium acetate + 50 mM TES + 1nooo ATCC-TMS solidified with 1.5% agar. Single colonies were picked from plates into SOB cultures. adding 50 µg/ml kanamycin in the case of C. *necator* isolates. and grown up overnight shaking at 30^°^C. Saturated cultures were submitted for whole genome sequencing (Angstrom Innovation) and cryo-preserved at -80°C after adding glycerol to 25% total volume. Whole genome sequencing was analyzed with breseq^44^ to identify mutations in each isolate compared to the starting ALE parent. To pick the top isolates for further testing in flasks. we measured growth curves across a range of concentrations for DMM and acetate, similar to assays performed above on evolved populations. and selected strains that reached the highest maximum OD_600_ at the highest concentrations of DMM and acetate across 72 hours of growth.

### Fed-batch Growth And PHA Production

Our top ALE isolates were assessed for growth in flasks with DMM and acetate, using a fed-batch process with acetic acid spike-ins to more closely represent a bioprocess on Mars. These assays also used our recently-developed MTM^15^ rather than ATCC-TMS to be more representative of trace metals present in leached martian regolith.

For C. *necator*, isolates were seeded from glycerol stocks into 50 ml SOB + 50 µg/ml kanamycin and grown overnight shaking at 30^°^C. Saturated cultures were then pelleted and normalized in TES-buffered saline to OD_600_ 2. TES buffer and kanamycin selection were used as laboratory conveniences for pH stability and contamination prevention, respectively; neither would be required in a Mars bioreactor. where pH would be managed by controlled acetic acid addition and contamination risk is inherently lower in a closed system. Sterile baffled flasks were filled with 100 ml of 15x DMM + 1.75% potassium acetate + 50 mM TES + lx MTM. and each flask was inoculated with 2 ml of OD normalized culture. Cultures were incubated shaking at 30^°^C for 72 hours. At 24. 48. and 72 hours. OD6oo was measured for each flask. and pH was measured with pH strips. At 48 hours when pH in cultures had reached pH 9. 800 µL of 20% acetic acid and 5.24 ml of 20% potassium acetate were added to each culture to reduce pH to 8 and replenish carbon for a total concentration of 1.25% acetate added back to each flask. To determine the volume of acetic acid needed to bring pH down to 8, we performed acetic acid titration on an additional culture, by adding 20% acetic acid in 100 µL steps and recording the resulting pH.

PHA was isolated and quantified following established methods^36^. Briefly, C. *necator* cultures were harvested at 72 hours by centrifugation. Pellets were then resuspended in 3 ml of 8% NaOCI and incubated with slow shaking at room temperature for two hours. dissolving non-bioplastic material. After incubation, bleached pellets were centrifuged, washed twice with 20 ml of deionized water. and dried overnight at 35^°^C. After drying, final PHA mass was measured and recorded (**Figure 7**).

**Figure 7.**
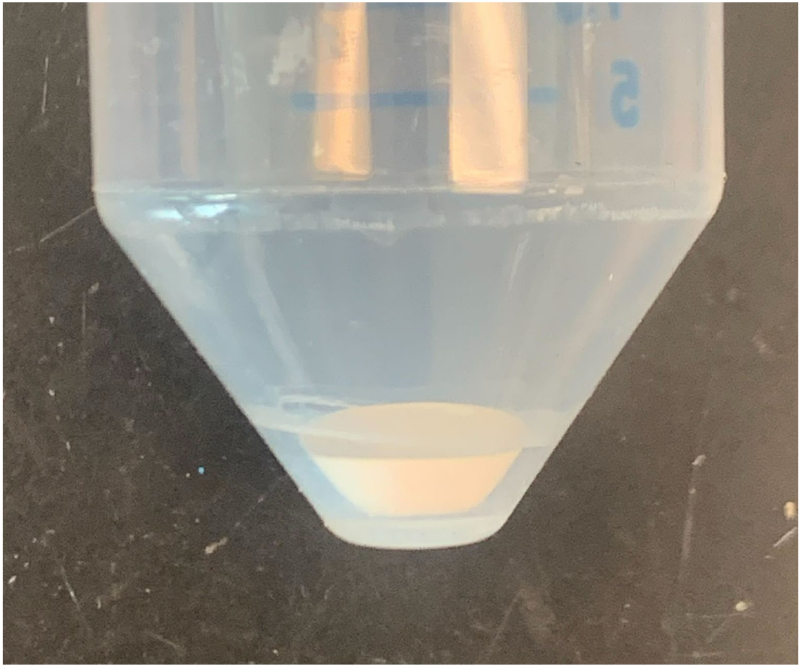
PHB pellet recovered after growth. Dried PHB from 100 ml *C. necator* grown in 15x DMM + 1.75% potassium acetate + 50mM TES + lx MTM.

For *P. putida*. we similarly characterized flask growth in Mars media, but did not pursue PHA purification. *P. putida* was evaluated identically to C. *necator*, but instead using a more challenging condition: 20x DMM + 2.5% potassium acetate + 50 mM TES + lx MTM. and seeded with 500 µL of culture at OD_600_ 2. pH and OD_600_ were measured at 24, 48, and 72 hours. and cultures were harvested at 72 hours. Unlike C. *necator. P. putida* cultures were not feed acetate and acetic acid because *P. putida* cultures had not increased in pH by 48 hours. *P. putida* cultures experienced greater lag due to more challenging conditions.

## ACKNOWLEDGEMENTS

This research was funded in part by The Astera Institute. In compliance with Astera’s Open Science policy^45^, this paper will not be submitted to a journal and is presented here in its final form. We thank Olesia Bushkova and Rachel Sevey for their assistance with scientific communication. We thank Hasan Celik at the Pines Magnetic Resonance Center’s Core NMR Facility at the University of California, Berkeley for NMR guidance and data acquisition.

